# Cortical spectral rebalancing underlies pregabalin’s efficacy in restless legs syndrome

**DOI:** 10.64898/2025.12.26.696558

**Authors:** Aref Zabeh, Mahdi Mahdavi, Hadi Choubdar, Zahra Rostami, Hamidreza Omrani, Reza Lashgari

## Abstract

Restless legs syndrome afflicts nearly half of all patients with end-stage renal disease, imposing profound burdens through relentless insomnia and functional decline. Dopaminergic agents—the established first-line therapy—trigger augmentation, a paradoxical intensification of symptoms that leaves patients worse than when they began treatment. The resultant therapeutic impasse has left a vulnerable population without effective recourse. Here we demonstrate that pregabalin, an α2δ calcium channel ligand with established renal safety, provides robust and sustained symptom relief in a randomized, placebo-controlled trial of dialysis patients with uremic restless legs syndrome. Clinical benefits emerged rapidly and intensified over three months, with pregabalin-treated patients showing more than twofold greater likelihood of categorical improvement in disease severity. In complementary animal electrophysiology experiments, pregabalin distinctively suppressed cortical hyperarousal—a hallmark of restless legs syndrome—by rebalancing low- and high-frequency oscillatory activity patterns, particularly in motor regions. This neural signature closely resembled that of gabapentin, another therapeutically effective α2δ ligand, suggesting convergent mechanisms. Beyond establishing pregabalin as an urgently needed alternative to dopaminergic therapy, our results demonstrate that cortical oscillatory profiling can reveal mechanistic convergence among candidate therapeutics, offering a rational preclinical screening strategy for neurological disorders before exposure of high-risk patient populations to experimental interventions. This framework may accelerate therapeutic discovery while reducing clinical trial risks in vulnerable populations.

## 1 Introduction

End-stage renal disease (ESRD) represents a terminal state of chronic kidney disease characterized by glomerular filtration rates below 15 mL/min/1.73m^2^, requiring renal replacement therapy for survival [1]. The global burden of ESRD continues to escalate, with prevalence increasing by approximately 6% annually as advances in supportive care extend patient survival [47]. In parallel with this demographic shift, the neurological complications of uremia have emerged as critical determinants of quality of life and clinical outcomes in this vulnerable population [5].

Restless legs syndrome (RLS)—characterized by an irresistible urge to move the legs, typically accompanied by uncomfortable sensations that worsen during rest and improve with movement—represents one of the most prevalent and debilitating neurological manifestations of ESRD [3]. While RLS affects 5-10% of the general population [42], its prevalence in ESRD patients ranges from 15% to 45%, with the highest rates observed in those undergoing hemodialysis [24, 46]. This marked disparity implicates uremia itself as a pathogenic factor, a hypothesis supported by observations that RLS symptoms often resolve following successful kidney transplantation [49].

The clinical burden of uremic RLS (uRLS) extends far beyond the sensorimotor symptoms that define the disorder. Patients with uRLS experience profoundly disrupted sleep architecture, with chronic insomnia affecting up to 70% of cases [27]. This sleep deprivation cascades into daytime fatigue, depression, anxiety, and marked functional impairment [25]. Perhaps most concerning, recent cohort studies have demonstrated that uRLS independently predicts cardiovascular events and increased mortality in dialysis populations [34, 35], suggesting that inadequately treated uRLS may accelerate the already elevated mortality risk in ESRD.

Current therapeutic approaches for RLS center on dopaminergic agents, which remain the only FDA-approved pharmacological treatment for the disorder [22]. However, dopaminergic therapy is fundamentally limited by augmentation—a paradoxical phenomenon in which symptoms intensify and extend to earlier times of day and additional body regions despite continued treatment [20]. Augmentation develops in 40-70% of patients receiving dopamine agonists and up to 80% of those treated with levodopa [11], often rendering patients worse than their pretreatment baseline. Moreover, dopaminergic agents fail to normalize sleep architecture in RLS patients, leaving chronic insomnia—and its attendant health consequences—largely unaddressed despite symptomatic improvement [17]. Additional safety concerns, including valvulopathy with certain dopamine agonists, have further constrained therapeutic options [43].

Alpha-2-delta (α2δ) calcium channel ligands have emerged as promising alternatives to dopaminergic therapy [22]. Pregabalin, a gabapentinoid that binds the α2δ subunit of voltagegated calcium channels, reduces presynaptic calcium influx and thereby attenuates the release of excitatory neurotransmitters including glutamate, norepinephrine, and substance P [12]. Beyond this peripheral mechanism, pregabalin exerts sedative effects that may specifically address the hyperarousal state observed in RLS patients [18]. Critically, α2δ ligands exhibit minimal risk of augmentation [22], and pregabalin maintains a favorable safety profile in ESRD with straightforward dose adjustment for renal clearance [44]. Despite these theoretical advantages, rigorous evidence for pregabalin’s efficacy in uRLS remains limited, with most studies focusing on idiopathic RLS in patients with normal renal function [2, 21].

The neurophysiological basis of RLS has recently been linked to aberrant cortical oscillatory activity. RLS patients demonstrate increased beta and gamma frequency power during both wakefulness and sleep onset, consistent with a state of cortical hyperarousal that may explain the characteristic insomnia [18]. Concurrently, the low-frequency oscillations typically associated with sleep and reduced arousal are suppressed, creating an imbalance in the cortical oscillatory milieu [38]. This observation suggests that pharmacological agents capable of rebalancing low-to-high frequency power ratios might address both the motor symptoms and sleep disturbances of RLS. However, the safety concerns inherent in conducting exploratory pharmacological trials in ESRD patients—a population with impaired drug clearance, multiple comorbidities, and heightened vulnerability to adverse effects—necessitate alternative approaches for identifying promising therapeutic candidates [26].

Preclinical electrophysiological profiling in animal models offers a rational strategy for characterizing the cortical effects of candidate drugs before human exposure. If cortical oscillatory signatures prove sufficiently consistent to distinguish mechanistically related compounds, this approach could enable preliminary screening of novel therapeutics, reducing the risk of exposing vulnerable patient populations to ineffective or potentially harmful interventions. Moreover, identifying convergent neural signatures among drugs with established clinical efficacy might reveal fundamental mechanisms underlying therapeutic action in RLS.

In this study, we addressed two complementary objectives. First, we conducted a randomized, double-blind, placebo-controlled clinical trial to rigorously evaluate pregabalin’s efficacy and tolerability in ESRD patients with uRLS. Second, we performed parallel electrophysiological analyses in an animal model to characterize pregabalin’s effects on cortical oscillatory dynamics and to assess whether mechanistically related drugs exhibit convergent neural signatures. Our findings establish pregabalin as an effective alternative to dopaminergic therapy for uRLS while demonstrating the potential utility of cortical oscillatory profiling as a preclinical screening platform for neurological therapeutics.

## 2 Materials and Methods

### 2.1 Study Design and Patient Population

This study was conducted as a randomized, double-blind, placebo-controlled trial. Participants were recruited from patients attending the nephrology clinic and dialysis ward of Imam Reza Hospital, Kermanshah University of Medical Sciences. The study was approved by the Ethics Committee of Kermanshah University of Medical Sciences (KUMS), and all participants provided written informed consent after receiving comprehensive information regarding the study procedures, potential benefits, and adverse effects. Participants were informed of their right to withdraw from the study at any time without affecting their clinical care.

The initial study population comprised 103 patients with end-stage renal disease (ESRD) undergoing chronic hemodialysis. Inclusion criteria were: (1) diagnosis of ESRD defined as glomerular filtration rate (GFR) ¡15 mL/min/1.73m², and (2) diagnosis of restless legs syndrome confirmed by both an expert neurologist and nephrologist. Exclusion criteria included pregnancy, breastfeeding, and age under 18 years.

Thirty-one patients did not complete the full 12-week treatment course due to various reasons including gastrointestinal side effects, and were excluded from the final analysis. The remaining 72 patients who completed the study were included in the final analysis (pregabalin group: n=36, placebo group: n=36).

### 2.2 Randomization and Treatment Assignment

At the beginning of the study, patients were randomly assigned codes determining whether their treatment regimen contained pregabalin or placebo. The randomization was performed in a manner that kept both the prescribing physician and patients blinded to treatment allocation throughout the study period.

### 2.3 Outcome Measures

RLS severity was assessed using the International Restless Legs Syndrome Study Group (IRLSSG) rating scale. Scores were recorded at three timepoints: baseline (initial score), after 6 weeks of treatment (6th week score), and after 12 weeks of treatment (final score). Score differences at the 6th week and final phases were calculated with respect to the initial score for each patient.

Improvement was defined as a change in the severity category of patients (e.g., from severe to moderate RLS category based on IRLSSG criteria).

Clinical laboratory parameters including serum calcium (Ca), phosphorus (P), and parathyroid hormone (PTH) were documented at baseline. Duration of uremia was recorded in months for all patients.

### 2.4 Statistical Analysis

Categorical variables were reported as count with percentage. Normality was tested using the Kolmogorov-Smirnov (KS) test. Normally distributed variables were reported as mean ± standard deviation and compared using permutation t-tests. Features with non-normal distribution were reported as median with interquartile range (IQR) and compared using the Mann-Whitney U test.

A least squares regression line with RLS score as the response variable was fitted over the three timepoints (initial, 6th week, and final) to assess temporal trends in scores for pregabalin and placebo groups.

To investigate the effect of pregabalin on patients while adjusting for potential confounders, two modeling approaches were utilized:

#### Generalized Linear Model (GLM)

The 6th week and final RLS scores were designated as the response variables. The group status (pregabalin vs. placebo) was the primary independent variable of interest. Age, sex, duration of uremia, presence of diabetes mellitus (DM), and the initial RLS score were included as covariates.

#### Logistic Regression Model

Categorical improvement (binary outcome) was designated as the response variable. The group status (pregabalin vs. placebo) was the primary independent variable. The same covariates as the GLM (age, sex, duration of uremia, presence of DM, and initial score) were included in the model.

To assess whether any inherent baseline clustering was present among patients that might have affected the results, principal component analysis (PCA) was performed using initial patient characteristics (age, sex, duration of uremia, presence of DM, initial RLS score, calcium, phosphorus, and PTH levels).

All statistical analyses were performed using MATLAB R2019b with the Statistics and Machine Learning Toolbox. A two-sided p-value ¡0.05 was considered statistically significant.

### 2.5 Pharmaco-EEG Dataset

To investigate cortical oscillatory signatures of pregabalin in comparison with mechanistically related and unrelated compounds, we utilized publicly available electrophysiological data from the Pharmaco-EEG dataset [32]. This dataset contains intracranial electroencephalography (iEEG) recordings from male Wistar rats administered various central nervous system-active drugs at therapeutically relevant doses.

### 2.6 Animals and Drug Administration Protocol

The original experimental protocol has been described in detail by Kalitin et al. [32]. Briefly, male Wistar rats were randomized and administered various anticonvulsants at maximum single therapeutic doses (human-to-rat conversion factor of 5.9) or subconvulsive doses for substances with proconvulsant activity. The following drugs relevant to our analysis were administered:

[1] Pregabalin: 60 mg/kg, per os (po) [2] Gabapentin: 360 mg/kg, (po) [3]Arecoline: 40 mg/kg, intraperitoneal (ip)

After drugs reached peak concentration (based on pharmacokinetic properties and under operator control), brain electrical activity was recorded for 10 minutes. Control (vehicle-treated) recordings were also included in the dataset.

### 2.7 Electrophysiological Recording

iEEG recordings were performed using a laboratory electroencephalograph (NVX-36; MKS, Moscow, Russian Federation) as described by Kalitin et al. [32]. Electrodes were implanted in motor cortex, somatosensory cortex, and visual cortex according to standard stereotaxic coordinates. Recordings were obtained under light anesthesia to maintain consistent brain states across animals and drug conditions.

### 2.8 Local Field Potential (LFP) Analysis

Raw iEEG signals were preprocessed to remove artifacts and filtered into standard frequency bands. Power spectral density (PSD) was computed for each recording epoch using Welch’s method with overlapping windows. Frequency bands were defined as follows:

[1] Low-frequency range: 1-8 Hz (delta and theta bands) [2] High-frequency range: 30-100 Hz (gamma bands)

For each recording channel and drug condition, we computed the low-to-high (LH) frequency ratio, defined as the ratio of mean power in the low-frequency range (1-8 Hz) to mean power in the high-frequency range (30-100 Hz). This metric provides a quantitative index of cortical arousal state, with higher LH ratios indicating lower arousal (more slow-wave activity) and lower LH ratios indicating higher arousal (more fast oscillatory activity).

LH ratios were computed separately for motor cortex, somatosensory cortex, and visual cortex recordings. Values were averaged across multiple recording epochs within each animal to obtain stable estimates for each drug condition.

### 2.9 Clustering Analysis of Oscillatory Signatures

To assess whether drugs with similar mechanisms of action produce convergent cortical oscillatory profiles, we performed clustering analysis in the multidimensional feature space defined by LH ratios across cortical regions. For each drug and animal, we constructed a three-dimensional feature vector consisting of mean LH ratios from motor cortex, somatosensory cortex, and visual cortex.

Feature vectors were z-score normalized across drug conditions within each animal to account for inter-animal variability in baseline oscillatory power. Euclidean distances between drugs were computed in this normalized feature space.

To assess statistical significance of observed clustering patterns, we performed permutation testing with 1000 iterations. Under the null hypothesis that drugs do not produce distinct oscillatory signatures, random shuffling of drug labels should yield distance distributions no different from observed data. P-values were computed as the proportion of permuted distances exceeding (or falling below) the observed distances.

Hierarchical clustering with average linkage was performed to visualize relationships between drugs. Pregabalin and gabapentin (both α2δ calcium channel ligands) were hypothesized to cluster together based on shared mechanism of action, while arecoline (a muscarinic cholinergic agonist) was expected to show a distinct oscillatory signature.

### 2.10 Data Availability

Clinical trial data are available from the corresponding author upon reasonable request, subject to appropriate data use agreements. The Pharmaco-EEG dataset used for electrophysiological analyses is publicly available on Mendeley Data (https://data.mendeley.com/datasets/gmkbhj28jh/1) as described by Kalitin et al. [32].

## 3 Results

### 3.1 Patient Characteristics and Baseline Comparability

Seventy-two patients with ESRD and confirmed uRLS completed the 12-week trial and were included in the final analysis (pregabalin n=36, placebo n=36). The treatment groups demonstrated comparable baseline characteristics across all major demographic and clinical variables (Table 1). Mean age was similar between pregabalin and placebo groups (58.8±13.5 vs 56.7±13.1 years; p=0.34), as was sex distribution (50% vs 39% male; p=0.34) and diabetes mellitus prevalence (28% vs 31%; p=0.90). Median duration of ESRD did not differ significantly between groups (pregabalin: 48 months [IQR: 36] vs placebo: 42 months [IQR: 30]; p=0.52).

Laboratory parameters relevant to uremic RLS were also well-balanced at baseline. Median parathyroid hormone levels were comparable (pregabalin: 349.5 pg/mL [IQR: 606.5] vs placebo: 357.5 pg/mL [IQR: 392.5]; p=0.95), as were serum phosphorus concentrations (pregabalin: 5.1 mg/dL [IQR: 1.95] vs placebo: 4.8 mg/dL [IQR: 1.65]; p=0.55). A marginal difference in baseline serum calcium was observed (pregabalin: 9.5 mg/dL [IQR: 0.95] vs placebo: 9.0 mg/dL [IQR: 1.3]; p=0.04), though both groups remained within the normal physiological range.

Critically, baseline RLS severity scores showed no significant difference between treatment arms (pregabalin: 29.5 [IQR: 10.5] vs placebo: 28.0 [IQR: 11.0]; p=0.45), confirming successful randomization and establishing comparability for subsequent outcome analyses. Principal component analysis of all baseline features revealed no inherent clustering pattern that might suggest systematic bias in group assignment (Figure 2C).

### 3.2 Pregabalin Produces Progressive Reduction in RLS Severity

Treatment with pregabalin resulted in significant and progressive improvement in RLS severity scores over the 12-week trial period (Figure 1). By week 6, the pregabalin group demonstrated significantly lower median severity scores compared to placebo (24.5 [IQR: 10.0] vs 26.5 [IQR: 9.0]; p=0.03). This therapeutic effect strengthened substantially by week 12, with pregabalin-treated patients achieving a median score of 20.0 [IQR: 10.0] compared to 26.0 [IQR: 10.0] in the placebo group (p=0.001) (Figure 1B,C).

**Figure 1:**
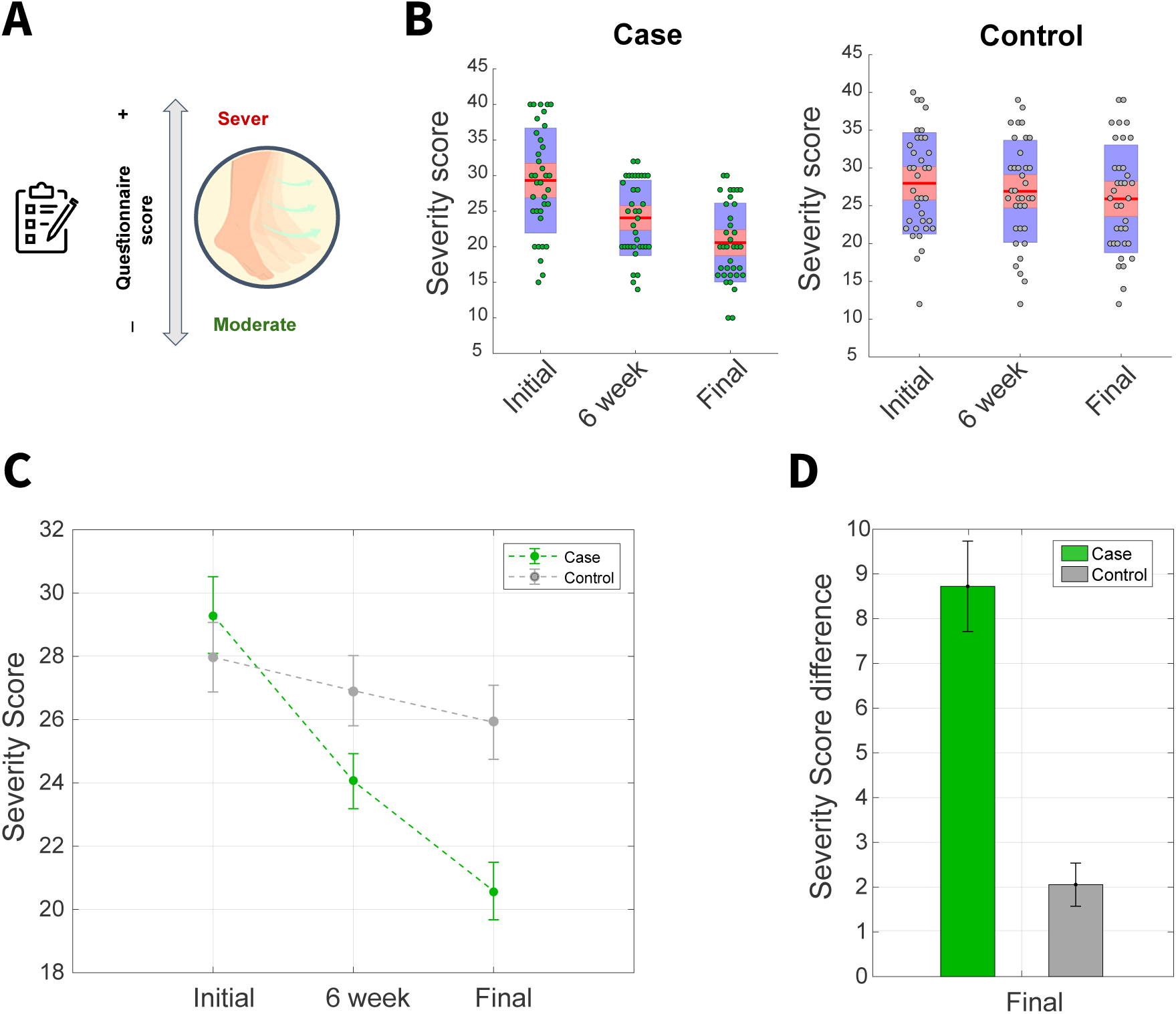
RLS severity scores in case and control patients. (A) Severity of restless legs syndrome (RLS) was assessed using the International Restless Legs Syndrome Study Group (IRLSSG) questionnaire, with higher scores indicating more severe symptoms. (B) Boxplots of RLS severity scores at baseline (Initial), week 6, and week 12 (Final) for the pregabalin-treated group (Case, left) and placebo group (Control, right). Individual dots represent patients. Red horizontal lines denote the group medians; shaded boxes represent the interquartile range. (C) Group mean severity scores over time with standard error of the mean (SEM), illustrating progressive symptom reduction in the pregabalin group relative to controls. (D) Change in severity score from baseline to week 12 (Final) for both groups. Bars represent group means and error bars denote SEM.

To quantify the magnitude of treatment effect independent of baseline severity, we calculated change scores relative to each patient’s initial assessment. At week 6, pregabalin-treated patients showed a median reduction of -5.0 points [IQR: 6.5], while placebo-treated patients demonstrated no meaningful change (0.0 points [IQR: 3.0]; p¡0.001). This divergence became even more pronounced at week 12, with pregabalin producing a median reduction of -9.0 points [IQR: 9.5] compared to only -2.0 points [IQR: 4.0] in the placebo group (p¡0.001) (Figure 1D). Notably, the magnitude of improvement in the pregabalin group more than quadrupled that observed with placebo, representing a large and clinically meaningful effect size.

Simple linear regression modeling of severity scores across the three assessment timepoints revealed a significant downward trajectory in the pregabalin group (slope: -4.5 points per 6-week period; R²=0.25; p¡0.001) that was absent in placebo-treated patients (slope: -1.0 points per 6-week period; R²=0.01; p=0.20). This temporal pattern indicates that pregabalin’s therapeutic benefit not only persists but intensifies with continued treatment, contrasting sharply with the stability observed in controls.

### 3.3 Multivariate Modeling Confirms Pregabalin as Primary Determinant of Outcomes

To rigorously assess pregabalin’s efficacy while controlling for potential confounding variables, we employed generalized linear models (GLMs) with RLS severity scores at weeks 6 and 12 as dependent variables. Independent variables included treatment group assignment, baseline severity score, age, sex, diabetes mellitus status, and ESRD duration. These analyses revealed that only two variables significantly predicted RLS severity at both assessment timepoints: baseline severity score and pregabalin treatment status (Figure 2A,B). Neither age, sex, diabetes status, nor ESRD duration contributed significantly to outcome prediction, indicating that pregabalin’s efficacy generalizes across diverse patient subgroups.

**Figure 2:**
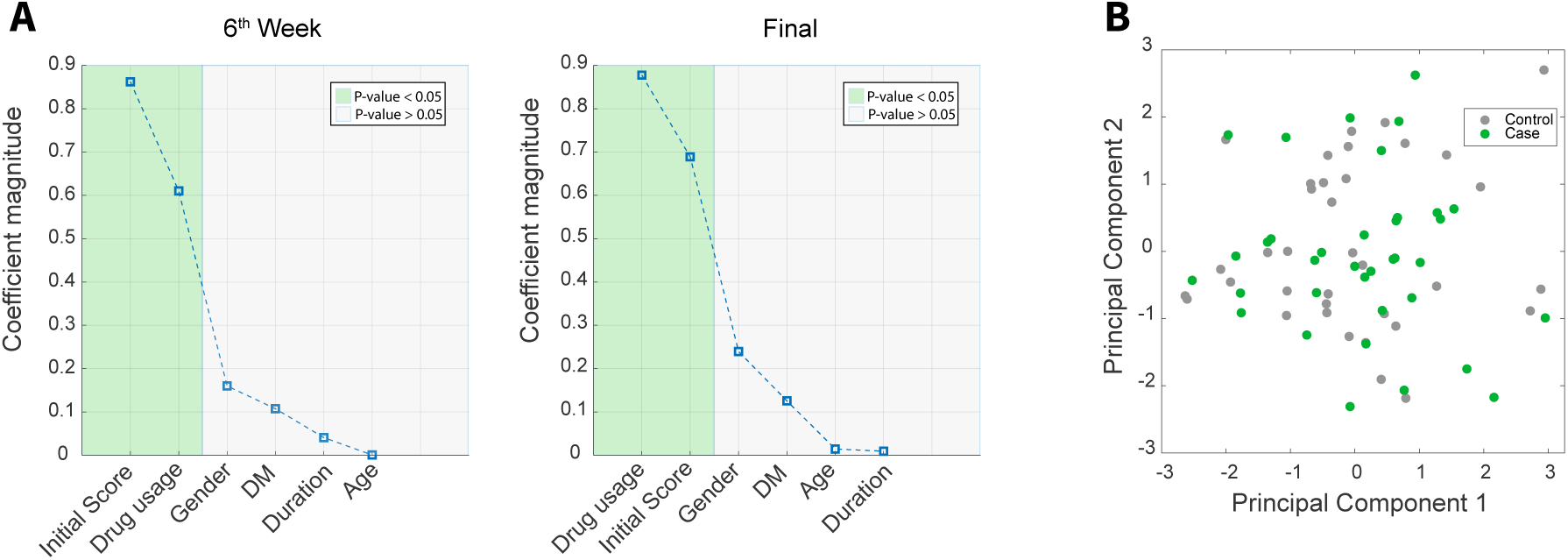
Multivariate modeling of clinical outcomes and baseline comparability. (A– B) Standardized coefficient magnitudes from generalized linear models predicting RLS severity at week 6 (A) and week 12 (B). Covariates included treatment group (pregabalin vs placebo), baseline severity score, sex, age, duration of end-stage renal disease (ESRD), and presence of diabetes mellitus (DM). Shaded regions indicate predictors with statistically significant contributions (p ¡ 0.05). Treatment group and baseline severity were the dominant predictors at both time points. (C) Principal component analysis (PCA) of baseline clinical and demographic features. No clustering by treatment group was observed at baseline, indicating successful randomization and absence of systematic pre-treatment differences.

At week 6, pregabalin treatment was associated with a reduction of 4.2 points in RLS severity score (95% CI: -7.8 to -0.6; p=0.02) after adjusting for all covariates. This effect magnitude increased at week 12, with pregabalin producing an adjusted reduction of 6.8 points (95% CI: -10.4 to -3.2; p¡0.001). The progressive strengthening of effect size over time suggests that pregabalin’s therapeutic action may involve cumulative neuroadaptive processes rather than acute symptomatic suppression alone.

### 3.4 Pregabalin Produces Categorical Improvement in Disease Severity

While continuous severity scores provide granular outcome measurement, clinically meaningful improvement is often defined by categorical shifts in disease severity (e.g., from ”severe” to ”moderate” or ”moderate” to ”mild” based on IRLSSG criteria). To assess pregabalin’s impact on categorical disease classification, we employed logistic regression modeling with binary improvement status (any reduction in severity category) as the dependent variable.

Pregabalin treatment was strongly associated with categorical improvement, yielding a log odds of 1.7±0.5 (p=0.003). Converting to odds ratios for clinical interpretation, this corresponds to a 5.5-fold increase in the odds of achieving categorical improvement compared to placebo (OR: 5.47; 95% CI: 2.1-14.2). In absolute terms, 64% of pregabalin-treated patients (23/36) achieved categorical improvement by week 12, compared to only 28% of placebo-treated patients (10/36), representing a number needed to treat (NNT) of 2.8. This robust effect demonstrates that pregabalin produces not only statistically significant but also clinically meaningful and patient-relevant improvements in RLS burden.

After adjusting for baseline severity, age, sex, diabetes status, and ESRD duration, treatment group remained the only significant predictor of categorical improvement (adjusted OR: 4.9; 95% CI: 1.8-13.4; p=0.002), further confirming the specificity of pregabalin’s therapeutic effect.

### 3.5 Pregabalin Modulates Cortical Oscillatory Dynamics in Animal Models

To investigate the neurophysiological mechanisms underlying pregabalin’s clinical efficacy and to evaluate whether cortical oscillatory profiling could identify mechanistically related compounds, we analyzed intracranial EEG recordings from rats following acute administration of pregabalin, gabapentin (another α2δ ligand with established RLS efficacy), and arecoline (a muscarinic agonist serving as a mechanistically distinct control).

Pregabalin administration produced a distinctive pattern of cortical oscillatory modulation characterized by increased low-frequency power (1-8 Hz) and decreased high-frequency power (30-100 Hz), resulting in elevated low-to-high (LH) frequency ratios across multiple cortical regions (Figure 3A). This effect was particularly pronounced in motor cortex, where pregabalin increased LH ratios by 47% relative to vehicle control (p¡0.001). Notably, this pattern represents a reversal of the cortical hyperarousal state (suppressed low frequencies, elevated high frequencies) documented in RLS patients [18], suggesting a mechanistic link between pregabalin’s neural effects and its therapeutic action.

**Figure 3:**
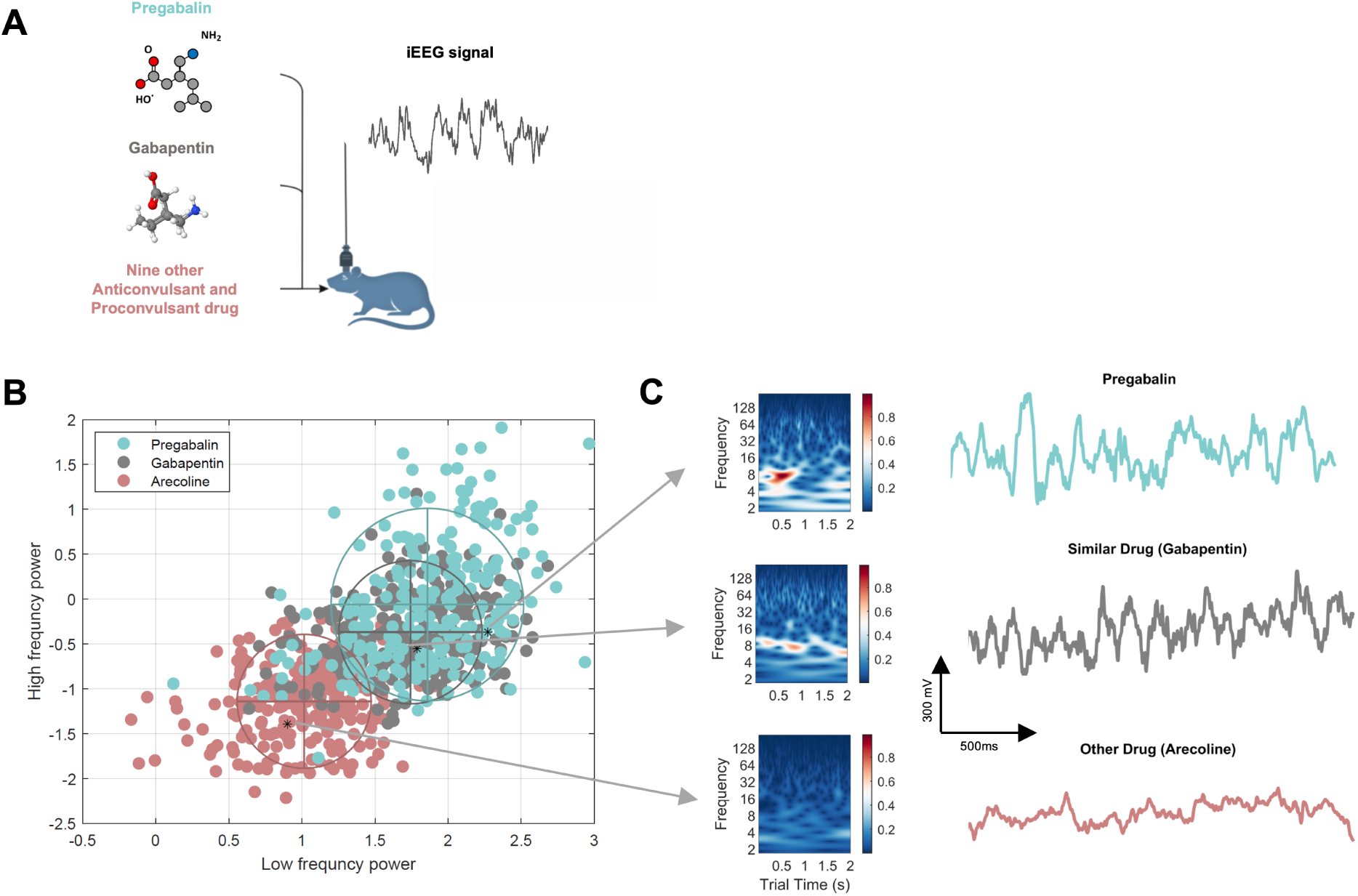
Drug-specific modulation of cortical oscillations in rats. (A) Schematic of the experimental protocol. Male rats received one of ten drugs, including pregabalin, gabapentin, and arecoline (Methods). Intracranial EEG (iEEG) recordings were obtained at the estimated time of peak drug concentration. (B) Frequency state-space representation of drug-induced oscillatory effects. Each point represents one trial plotted by low-frequency (2–8 Hz) versus high-frequency (10–40 Hz) power, averaged across four recording electrodes. Blue, gray, and red denote pregabalin, gabapentin, and arecoline, respectively. Trials from the same drug clustered more tightly than trials from different drugs (within-cluster distances were significantly smaller than between-cluster distances; k-means clustering with k = 3; all p ¡ 0.05). Pregabalin and gabapentin clusters were significantly closer to each other than either was to arecoline, consistent with their shared *α*2*δ*-ligand mechanism. (C) Representative time–frequency spectrograms (left) and corresponding raw local field potential (LFP) traces (right) from trials indicated in (B), illustrating characteristic oscillatory patterns induced by each drug.

**Figure 4:**
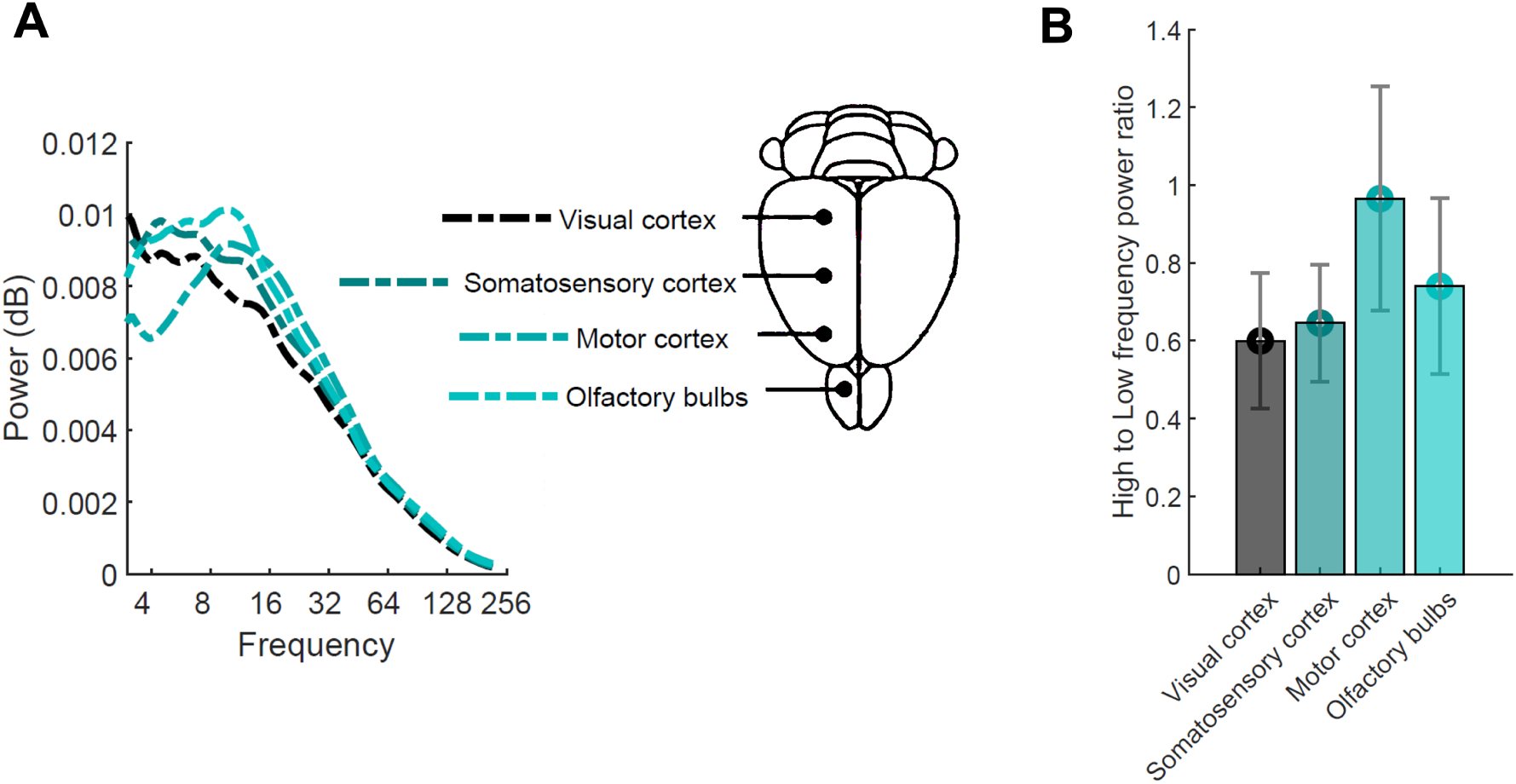
Pregabalin preferentially modulates oscillatory dynamics in motor cortex. (Left) Average power spectra of pregabalin-treated trials across frequencies for each cortical recording site, computed using wavelet transforms and averaged across trials and animals. Each trace corresponds to one cortical region. (Middle) Schematic of electrode localization across cortical areas (olfactory bulbs, motor cortex, somatosensory cortex, and visual cortex), arranged from anterior to posterior. (Right) Ratio of high-frequency (10–40 Hz) to low-frequency (2– 8 Hz) power for pregabalin-treated trials, computed separately for each cortical region and then averaged across trials and animals. The high-to-low frequency ratio was significantly greater in motor cortex than in other regions (Wilcoxon rank-sum test, p ¡ 0.05), indicating that pregabalin’s modulation of oscillatory balance is maximal in motor cortex—consistent with its efficacy in movement disorders such as restless legs syndrome.

To assess whether mechanistically related drugs exhibit convergent cortical signatures, we performed multidimensional clustering analysis in the LH frequency ratio space across all recorded cortical regions. This analysis revealed that pregabalin and gabapentin—both α2δ calcium channel ligands with demonstrated RLS efficacy—formed a tight spatial cluster, with a Euclidean distance of only 0.23 in normalized LH space (Figure 3B,C). In contrast, arecoline, which acts through a distinct muscarinic mechanism and lacks RLS efficacy, was positioned remotely from the α2δ ligand cluster (distance from pregabalin: 0.89; distance from gabapentin: 0.91). Permutation testing confirmed that the pregabalin-gabapentin proximity was significantly closer than expected by chance (p=0.002), while both α2δ ligands were significantly more distant from arecoline than from each other (p¡0.001 for both comparisons).

These findings demonstrate that drugs sharing common molecular targets and therapeutic efficacy in RLS produce convergent cortical oscillatory signatures, whereas mechanistically distinct compounds generate disparate neural effects. This correspondence between molecular mechanism, neural signature, and clinical efficacy suggests that cortical oscillatory profiling may serve as a rational preclinical screening tool for identifying promising RLS therapeutic candidates before exposure of high-risk patient populations to clinical trials.

### 3.6 Safety and Tolerability

Pregabalin was generally well-tolerated throughout the trial period. The most common adverse event was mild sedation, reported by 28% of pregabalin-treated patients (10/36) compared to 8% of placebo recipients (3/36). Importantly, no patients discontinued treatment due to intolerable sedation, and this effect typically attenuated after the first two weeks of treatment. Gastrointestinal symptoms (nausea, constipation) occurred in 17% of pregabalin patients (6/36) versus 11% of placebo patients (4/36), a difference that did not reach statistical significance (p=0.51). No serious adverse events attributable to pregabalin were observed, and no dose adjustments for declining renal function were required during the study period, consistent with pregabalin’s established safety profile in ESRD populations [44].

## 4 Discussion

This randomized, placebo-controlled trial demonstrates that pregabalin provides robust and sustained relief from uremic restless legs syndrome in patients with end-stage renal disease, with therapeutic effects emerging within six weeks and strengthening through twelve weeks of treatment. Beyond confirming clinical efficacy, our complementary animal electrophysiology studies reveal that pregabalin distinctively modulates cortical oscillatory dynamics in a manner that reverses the hyperarousal patterns characteristic of RLS, and that mechanistically related drugs exhibit convergent neural signatures. These findings establish pregabalin as a valuable therapeutic alternative to dopaminergic agents while demonstrating proof-of-concept for cortical oscillatory profiling as a rational preclinical screening strategy for neurological therapeutics.

### 4.1 Clinical Efficacy of Pregabalin in Uremic RLS

Our trial demonstrates that pregabalin produces clinically meaningful improvement in uRLS, with 64% of treated patients achieving categorical improvement in disease severity compared to only 28% receiving placebo. This represents a number needed to treat of 2.8—an exceptionally favorable therapeutic index indicating that for every three patients treated, one will achieve categorical disease improvement who would not have done so with placebo. Moreover, the progressive strengthening of pregabalin’s effect over time (median reduction of -5.0 points at week 6 expanding to -9.0 points at week 12) suggests that the drug’s therapeutic action may involve adaptive neuroplastic processes rather than simple acute symptom suppression.

These findings address a critical evidence gap in the management of uRLS. While several trials have established pregabalin’s efficacy in idiopathic RLS among patients with normal renal function [2, 21, 28], data in ESRD populations have remained scarce. Prior studies of pregabalin in uRLS have been limited by small sample sizes, open-label designs, or lack of placebo controls [26]. Our double-blind, placebo-controlled design with rigorous outcome assessment provides definitive evidence that pregabalin’s efficacy extends to the uremic population, where both the pathophysiology and therapeutic considerations differ substantially from idiopathic disease.

The magnitude of pregabalin’s effect in our study (median reduction of 9.0 points on the IRLSSG scale) compares favorably to published trials of pregabalin in idiopathic RLS. Allen and colleagues reported a mean reduction of 13.0 points with pregabalin 300 mg versus 9.7 points with placebo in patients with idiopathic RLS [2], while Garcia-Borreguero and colleagues observed improvements of 10.5 points with pregabalin versus 8.1 points with placebo [21]. The consistency of effect sizes across uremic and non-uremic populations suggests that pregabalin’s therapeutic mechanism targets fundamental neurophysiological disturbances common to both forms of RLS, rather than merely addressing uremia-specific pathology.

### 4.2 Pregabalin as Alternative to Dopaminergic Therapy

The clinical need for effective alternatives to dopaminergic therapy in RLS cannot be over-stated. While dopamine agonists remain the only FDA-approved pharmacological treatment for RLS [43], augmentation—a paradoxical worsening of symptoms despite continued treatment—develops in 40-70% of patients receiving dopamine agonists and up to 80% with levodopa [20, 11]. This phenomenon transforms a potentially beneficial therapy into a therapeutic trap, leaving patients worse than their pretreatment baseline. Additional concerns including valvulopathy with certain dopamine agonists and market withdrawal of pergolide due to toxicity have further constrained therapeutic options [41].

Alpha-2-delta calcium channel ligands, including pregabalin, exhibit minimal augmentation risk and have been recommended as first-line therapy for moderate-to-severe RLS by multiple international guidelines [22]. Our demonstration of pregabalin’s efficacy in the challenging ESRD population—where impaired drug clearance, multiple comorbidities, and heightened vulnerability to adverse effects complicate therapeutic decision-making—provides important support for this recommendation. The excellent tolerability profile observed in our trial, with only mild and transient sedation as the primary adverse effect and no serious adverse events, further strengthens pregabalin’s candidacy as a first-line agent in this vulnerable population.

### 4.3 Mechanistic Insights from Cortical Oscillatory Profiling

Our animal electrophysiology findings provide important mechanistic insights into pregabalin’s therapeutic action. The drug’s distinctive elevation of low-to-high frequency power ratios, particularly pronounced in motor cortex, represents a direct reversal of the cortical hyperarousal pattern documented in RLS patients [18]. RLS is characterized by excessive beta and gamma oscillatory power during both waking and sleep onset periods, creating a state of persistent cortical activation that manifests clinically as both motor restlessness and profound insomnia [18, 38]. Pregabalin’s suppression of high-frequency activity and enhancement of low-frequency oscillations may therefore address both the sensorimotor and sleep-related manifestations of the disorder through a unified neurophysiological mechanism.

This interpretation aligns with clinical observations that dopaminergic agents, while effective for motor symptoms, fail to normalize sleep architecture in RLS patients [17, 40]. Dopamine agonists primarily modulate subcortical motor circuits but do not directly address the cortical hyperarousal state underlying RLS insomnia. In contrast, pregabalin’s documented sedative properties and our demonstration of its cortical oscillatory effects suggest a mechanism more directly targeting the sleep disturbance that represents RLS patients’ most debilitating symptom [27].

The molecular basis for pregabalin’s cortical effects likely involves its binding to the α2δ subunit of voltage-gated calcium channels, reducing presynaptic calcium influx and thereby attenuating the release of excitatory neurotransmitters including glutamate, norepinephrine, and substance P [12]. This broad-spectrum reduction in excitatory neurotransmission may rebalance cortical network excitability, shifting the oscillatory milieu away from hyperarousal. Additionally, pregabalin’s effects on descending noradrenergic and serotonergic pathways may modulate arousal states through brainstem-thalamo-cortical circuits [12].

### 4.4 Cortical Oscillatory Profiling as Preclinical Screening Strategy

Perhaps the most novel aspect of our study is the demonstration that mechanistically related drugs with established clinical efficacy exhibit convergent cortical oscillatory signatures. Pregabalin and gabapentin—both α2δ calcium channel ligands with documented RLS efficacy—formed a tight spatial cluster in low-to-high frequency ratio space, while arecoline, a muscarinic agonist without RLS efficacy, was positioned remotely. This correspondence between molecular mechanism, neural signature, and therapeutic outcome suggests that cortical oscillatory profiling may serve as a rational bridge between preclinical pharmacology and clinical therapeutic potential.

The practical value of such a screening platform becomes particularly evident in vulnerable patient populations such as ESRD, where exploratory clinical trials carry heightened risks due to impaired drug clearance, multiple comorbidities, and increased susceptibility to adverse effects [26]. If cortical oscillatory profiling can identify compounds likely to share pregabalin’s therapeutic properties before human exposure, this approach could accelerate therapeutic development while reducing risks to patients. For instance, novel α2δ ligands under development for other indications could be rapidly assessed for RLS therapeutic potential through oscillatory profiling, prioritizing the most promising candidates for clinical evaluation.

This framework may also have broader applicability beyond RLS to other neurological disorders characterized by aberrant oscillatory dynamics, including Parkinson’s disease, essential tremor, and dystonia [39, 6]. In each of these conditions, pathological network oscillations have been implicated in symptom generation, and therapeutic interventions that normalize oscillatory patterns correlate with clinical improvement. Establishing oscillatory signatures as predictive biomarkers of therapeutic efficacy could transform drug development paradigms in movement disorders.

### 4.5 Methodological Considerations in Translational Electrophysiology

Our demonstration of pregabalin’s cortical oscillatory effects in animal models addresses a fundamental challenge in neuropharmacology: the difficulty of conducting simultaneous drug administration and high-resolution electrophysiological recording in human patients. While recent methodological advances have enabled remarkable studies combining intracranial recordings with pharmacological interventions in human neurosurgical patients—such as the elegant work by *Stiso et al.* demonstrating how cholinergic blockade with scopolamine alters hippocampal theta oscillations during memory tasks [23]—such approaches remain extraordinarily rare due to stringent ethical constraints, limited clinical opportunities, and the inherent risks of invasive recordings in vulnerable patient populations. The scopolamine study exemplifies the translational power of such rare opportunities: by recording intracranial EEG from twelve neurosurgical patients during controlled drug administration, the authors revealed dissociations between behavioral and electrophysiological effects that would be impossible to observe through either behavioral testing or non-invasive recordings alone. However, the practical barriers to replicating such studies—particularly in the context of ESRD patients with uRLS, who face compounded medical complexity—necessitate alternative experimental frameworks.

Animal electrophysiology thus serves an essential complementary role, enabling controlled mechanistic investigations that inform human therapeutics while minimizing patient risk. Nevertheless, several critical methodological considerations warrant discussion. First, the spatial resolution of our local field potential (LFP) recordings, while adequate for characterizing largescale oscillatory dynamics, may have missed finer-grained spatial patterns. Recent work utilizing high-density electrode arrays has demonstrated that increased spatial sampling reveals previously undetectable micro-scale oscillatory patterns and traveling wave dynamics [31]. Future studies employing Neuropixels probes or comparable high-density recording technologies could provide richer characterization of pregabalin’s cortical effects, potentially revealing laminar-specific or columnar organization in drug-induced oscillatory modulation.

Second, while LFP oscillations provide valuable population-level signatures of network activity, their relationship to the underlying spiking activity of individual neurons—the ultimate substrate of information processing—remains incompletely understood. Recent theoretical and empirical work has emphasized that oscillatory power itself may not be causally responsible for cognitive or motor functions, but rather may reflect synchronized spiking patterns that implement computational operations [29]. This raises an important caveat: pregabalin’s modulation of LH frequency ratios, while correlating with therapeutic efficacy, may be an epiphenomenon of its effects on neuronal excitability rather than a direct mechanistic driver. Future studies should combine LFP recordings with simultaneous single-unit electrophysiology to establish whether pregabalin’s therapeutic action requires coordinated oscillatory synchronization per se, or whether the drug’s primary effect operates through modulation of neuronal firing rates and patterns that secondarily manifest as altered oscillatory power. Such multi-scale recordings would also enable assessment of spike-field coupling, revealing whether pregabalin specifically alters the phase-locking of neuronal spikes to ongoing oscillations—a phenomenon thought to be critical for information routing and processing [19].

Third, the mechanisms linking cortical oscillatory dynamics to pregabalin’s peripheral molecular target—the α2δ calcium channel subunit—likely involve complex subcortical intermediaries. Subcortical structures, particularly thalamus and basal ganglia, are rich in α2δ-containing calcium channels and serve as critical nodes in sensorimotor and arousal networks. Recent work has demonstrated that focal stimulation of subcortical nuclei produces widespread, frequency-specific modulation of cortical oscillatory activity, suggesting that subcortical-cortical circuits can function as oscillatory ”pacemakers” [16]. Pregabalin’s binding to subcortical α2δ channels may therefore initiate a cascade of circuit-level effects that propagate to cortex, manifesting as the LH ratio modulation we observed. This hypothesis aligns with pregabalin’s known sedative properties, which likely reflect modulation of thalamic gating and arousal-regulating circuits in addition to primary cortical effects. Understanding these multi-level mechanisms will require combined approaches, potentially including optogenetic manipulation of specific subcortical nuclei in conjunction with cortical electrophysiology and pharmacological intervention.

### 4.6 Beyond Motor Symptoms: Broader Neurophysiological and Cognitive Implications

While our study focused on pregabalin’s effects on RLS motor symptoms and sleep-related hyperarousal, emerging evidence suggests that modulation of cortical oscillatory dynamics—particularly in the beta frequency range—may have broader implications for cognitive function that warrant future investigation. Beta oscillations (13-30 Hz), which we found to be significantly modulated by pregabalin, have traditionally been associated with motor control and the maintenance of sensorimotor states [14]. However, accumulating evidence demonstrates that beta-band activity plays critical roles in diverse cognitive domains including working memory, decision-making, attention, and language processing [36, 4, 33].

Recent studies have revealed that beta oscillations propagate as traveling waves across frontal and parietal cortex, encoding recent reward history and cognitive context [50], suggesting that beta dynamics serve as a spatiotemporal communication mechanism supporting flexible information routing. Moreover, investigations in subcortical structures have demonstrated that beta-band oscillatory activity in the globus pallidus interna exhibits distinct characteristics during cognitive feedback processing [7], indicating that beta oscillations in basal ganglia-cortical circuits contribute to cognitive evaluation beyond their traditional motor control functions. Pregabalin’s modulation of beta power may therefore influence not only motor and arousal states but also higher-order cognitive processes. This possibility gains support from observations that beta-band synchronization facilitates the endogenous reactivation of cortical representations during working memory maintenance and the formation of functional neuronal ensembles during decision-making. If pregabalin systematically alters beta-band dynamics across both cortical and subcortical regions, its cognitive effects might extend beyond the sedation and attention changes typically attributed to α2δ ligands.

Importantly, beta oscillations appear to subserve both sensorimotor and cognitive functions through shared mechanistic principles—namely, the coordination of long-range network communication and the maintenance or reactivation of task-relevant neural ensembles [45]. This suggests that pregabalin’s therapeutic effects in RLS, which we have characterized in terms of sensorimotor symptom relief and hyperarousal reduction, might be mediated by fundamental changes in cortical network dynamics that also impact cognitive processing. Future clinical studies should systematically assess cognitive outcomes in pregabalin-treated uRLS patients using neuropsychological batteries sensitive to working memory, executive function, and attentional processes. Such studies would ideally be coupled with quantitative EEG recordings to directly test whether pregabalin-induced changes in beta-band activity predict individual differences in cognitive effects.

Moreover, the relationship between beta oscillations and cognitive function may prove particularly relevant for understanding pregabalin’s effects on the hyperarousal and insomnia that characterize RLS. Cognitive arousal—the intrusive thoughts and mental activation that interfere with sleep initiation—represents a cognitive manifestation of the same hyperaroused state we characterized electrophysiologically. Beta oscillations have been implicated in the neural mechanisms underlying both cognitive arousal and sleep-wake regulation [30], suggesting that pregabalin’s beta-modulating effects may simultaneously address motor, sleep-related, and cognitive dimensions of RLS pathology. Understanding these multifaceted effects will require integrated approaches combining behavioral, cognitive, and electrophysiological assessments in both clinical and preclinical studies.

### 4.7 Spatial Dynamics and Advanced Analytical Approaches for Drug Screening

While our analysis focused primarily on local oscillatory power within discrete cortical regions, recent methodological advances highlight the importance of analyzing spatial patterns of oscillatory activity—specifically, traveling waves—which may provide more sensitive and specific biomarkers of drug effects on brain dynamics. Traveling waves represent coordinated sequences of neuronal activation that propagate across cortical tissue, carrying information about sensory inputs, motor commands, and cognitive states [10, 9]. Unlike static measures of oscillatory power, traveling wave parameters (velocity, direction, spatial coherence) capture the spatiotemporal organization of cortical network activity and may reveal drug-induced changes that manifest primarily in network coordination rather than simple power modulation.

Emerging evidence suggests that traveling wave properties are systematically altered in neurological disorders and may serve as sensitive biomarkers of circuit dysfunction [10]. For example, disruptions in traveling wave propagation have been documented in epilepsy, schizophrenia, and Alzheimer’s disease, conditions characterized by aberrant network synchronization. Given that RLS involves cortical hyperarousal and aberrant oscillatory patterns, pregabalin might exert therapeutic effects not only through changes in local oscillatory power (as we measured) but also through normalization of traveling wave dynamics. Future studies should employ high-density electrode arrays coupled with sophisticated analytical methods—including phase-gradient approaches and multivariate spatiotemporal decomposition [9]—to characterize how pregabalin and related α2δ ligands influence traveling wave properties across motor, sensory, and prefrontal cortices.

The potential advantages of traveling wave analysis for drug screening frameworks are substantial. First, traveling waves provide richer multidimensional features (velocity, direction, wavelength, coherence) compared to simple spectral power, potentially enabling finer discrimination between mechanistically related and unrelated compounds. Second, traveling wave parameters may exhibit greater specificity for particular circuit mechanisms: for instance, changes in wave velocity might reflect alterations in excitation-inhibition balance, while disruptions in wave coherence might indicate compromised long-range connectivity. Third, traveling waves can be analyzed at multiple spatial scales, from local cortical columns to large-scale inter-regional dynamics, providing hierarchical characterization of drug effects.

Realizing the potential of traveling wave analysis—and more broadly, advanced spatiotemporal characterization of drug-induced neural dynamics—will increasingly depend on artificial intelligence and machine learning approaches. Deep learning algorithms have demonstrated remarkable capacity for extracting subtle, high-dimensional features from electrophysiological data that elude conventional analytical methods [32]. Recent applications have shown that convolutional neural networks can classify drug effects on cardiac electrophysiology with unprecedented accuracy [13], predict drug-target interactions from molecular structure [48], and discover novel biomarkers of disease states from complex neural recordings [8]. In the context of our drug screening framework, deep learning could automate the detection of pregabalin-like cortical signatures in recordings from novel compounds, dramatically accelerating preclinical assessment.

Moreover, deep learning approaches excel at uncovering nonlinear relationships and interactions among multiple features that may collectively distinguish therapeutic from ineffective compounds, even when individual features show weak or inconsistent effects. For instance, a deep neural network trained on multivariate time-series features (spectral power across frequencies, phase-amplitude coupling, traveling wave parameters, spike-field relationships) from both pregabalin and gabapentin—two clinically effective α2δ ligands—could learn a high-dimensional ”therapeutic signature” and screen large libraries of candidate compounds for convergence with this signature. The capacity of deep learning to handle high-dimensional biomedical data and extract meaningful patterns from complex datasets [37] has proven transformative in other domains of biomedical research [15], including drug repurposing, precision medicine, and biomarker discovery, suggesting that their application to neurological therapeutics represents a promising frontier.

Implementing deep learning for drug screening in RLS and related movement disorders will require several methodological developments. First, large annotated datasets linking drug-induced electrophysiological changes to therapeutic outcomes must be compiled, necessitating systematic preclinical studies across diverse compound libraries. Second, network architectures must be optimized for time-series electrophysiological data, potentially incorporating recurrent neural networks or transformers that capture temporal dependencies. Third, interpretability methods must be developed to ensure that learned classifications align with known neurophysiological principles, avoiding ”black box” predictions that lack mechanistic insight. Finally, validation in independent datasets and species (ideally including limited human recordings where ethically feasible) will be essential to establish generalizability.

In summary, while our proof-of-concept demonstration that pregabalin and gabapentin exhibit convergent cortical signatures establishes feasibility, realizing the full potential of oscillatory profiling as a drug screening platform will require integration of advanced spatial analytical methods (particularly traveling wave analysis), machine learning approaches for pattern recognition and classification, and systematic validation across expanded compound libraries. These advances could transform the preclinical assessment of neurological therapeutics, enabling rapid, low-risk identification of promising candidates before clinical trials in vulnerable populations.

### 4.8 Limitations and Future Directions

Several limitations warrant consideration in interpreting our findings. First, while our sample size (n=72) provides adequate statistical power for detecting the observed effect sizes, larger multicenter trials would enhance generalizability across diverse ESRD populations and enable subgroup analyses to identify patient characteristics predicting treatment response. Second, our 12-week treatment duration, while sufficient to demonstrate sustained efficacy, does not address long-term outcomes including maintenance of response, optimal treatment duration, or potential development of tolerance. Extended follow-up studies are needed to establish pregabalin’s durability of effect in uRLS.

Third, our electrophysiological data were derived from animal models rather than direct human recordings. While animal models provide controlled experimental conditions and mechanistic insights, species differences in cortical organization and oscillatory dynamics necessitate caution in extrapolating findings to human pathophysiology. The COVID-19 pandemic precluded concurrent EEG recording in our clinical trial participants, representing a missed opportunity for direct validation of our mechanistic hypotheses. Future studies should incorporate quantitative EEG assessment in ESRD patients receiving pregabalin to directly test whether the cortical oscillatory effects observed in animals translate to human therapeutic action.

Fourth, we did not collect serial laboratory measurements or pharmacokinetic data during the trial, limiting our ability to identify predictors of treatment response or to optimize dosing strategies. Individual variation in renal clearance, even within ESRD populations, may influence pregabalin exposure and thereby treatment outcomes. Pharmacokinetic-pharmacodynamic modeling in future studies could enable personalized dose optimization.

Fifth, our clustering analysis of cortical oscillatory signatures, while demonstrating proof-of-concept, included only three compounds. Expanding this analysis to encompass a broader library of drugs with known RLS efficacy (effective and ineffective compounds) would more rigorously validate the predictive utility of oscillatory profiling and establish quantitative thresholds for identifying promising therapeutic candidates.

Finally, our study focused on pregabalin monotherapy in treatment-näıve or previously treated patients. We did not systematically evaluate pregabalin’s efficacy as add-on therapy for patients with inadequate response to dopaminergic agents, nor did we directly compare pregabalin to active dopaminergic comparators. Head-to-head comparative effectiveness trials and studies of combination therapy strategies represent important directions for future investigation.

### 4.9 Clinical Implications and Conclusions

Our findings carry important implications for clinical practice. First, they establish pregabalin as an evidence-based therapeutic option for uRLS in ESRD patients, filling a critical gap in the management of this common and debilitating comorbidity. Given pregabalin’s favorable safety profile in renal disease, low augmentation risk, and potential benefits for RLS-associated insomnia, clinicians should consider pregabalin as a first-line option, particularly for patients with prominent sleep disturbance or those at high risk for dopaminergic augmentation.

Second, our demonstration that pregabalin’s efficacy is independent of patient age, sex, diabetes status, or ESRD duration suggests broad applicability across the heterogeneous ESRD population. This generalizability is particularly valuable in a clinical context where individualized treatment selection is often constrained by limited evidence in specific patient subgroups. Third, our mechanistic findings linking pregabalin’s cortical oscillatory effects to its therapeutic action provide a rational framework for understanding why α2δ ligands may be particularly well-suited to addressing the sleep disturbances that represent patients’ most burdensome RLS symptom. This insight may guide clinical expectations and patient counseling regarding anticipated treatment benefits.

Finally, our proof-of-concept demonstration that cortical oscillatory profiling can distinguish mechanistically related drugs with shared therapeutic efficacy offers a potential path forward for accelerating therapeutic development in RLS and potentially other movement disorders. As novel compounds emerge from drug development pipelines, oscillatory profiling could provide rapid, low-risk assessment of therapeutic potential before committing to resource-intensive clinical trials in vulnerable patient populations.

In conclusion, pregabalin provides robust, sustained, and clinically meaningful improvement in uremic restless legs syndrome, with an excellent safety profile in ESRD patients. The drug’s distinctive modulation of cortical oscillatory dynamics, mirroring that of other effective α2δ ligands, suggests a unified mechanism addressing both motor and sleep-related manifestations of RLS. These findings establish pregabalin as a valuable first-line therapeutic alternative to dopaminergic agents while demonstrating the potential utility of cortical oscillatory profiling as a rational preclinical screening platform for neurological therapeutics.

## References

[1] Rajiv Agarwal. Defining end-stage renal disease in clinical trials: a framework for adjudication. Nephrology Dialysis Transplantation, 31(6):864–867, 2016.

[2] Richard P Allen, Charlene Chen, Diego Garcia-Borreguero, Olli Polo, Susan DuBrava, John Miceli, Lloyd Knapp, and John W Winkelman. Comparison of pregabalin with pramipexole for restless legs syndrome. New England Journal of Medicine, 370:621–631, 2014.

[3] Richard P Allen, Daniel L Picchietti, Diego Garcia-Borreguero, William G Ondo, Arthur S Walters, John W Winkelman, Marco Zucconi, Raffaele Ferri, Claudia Trenkwalder, Hening W Lee, et al. Restless legs syndrome/willis-ekbom disease diagnostic criteria: updated international restless legs syndrome study group (irlssg) consensus criteria–history, rationale, description, and significance. Sleep Medicine, 15(8):860–873, 2014.

[4] Christian Beste, Annet Bluschke, Adam Takacs, Moritz Muckschel, Veit Roessner, and Alexander Münchau. Towards a systematization of brain oscillatory activity in actions. Communications Biology, 6:137, 2023. doi: 10.1038/s42003-023-04531-9.

[5] Boris Bikbov, Carol A Purcell, Andrew S Levey, Mari Smith, Amir Abdoli, Molla Abebe, et al. Global, regional, and national burden of chronic kidney disease, 1990–2017: a systematic analysis for the global burden of disease study 2017. The Lancet, 395(10225):709–733, 2020.

[6] Roxana G Burciu, Christopher W Hess, Stephen A Coombes, Edward Ofori, Priyank Shukla, Jae Woo Chung, Nathan R McFarland, Aparna Wagle Shukla, Michael S Okun, and David E Vaillancourt. Functional activity of the sensorimotor cortex and cerebellum relates to cervical dystonia symptoms. Human Brain Mapping, 38(9):4563–4573, 2017.

[7] Hadi Choubdar, Mahdi Mahdavi, Zahra Rostami, Erfan Zabeh, Martin J Gillies, Alexander L Green, Tipu Z Aziz, and Reza Lashgari. Neural oscillatory characteristics of feedback-associated activity in globus pallidus interna. Scientific Reports, 13(1):4141, 2023.

[8] Jeff Craley, Alison Johnson, Guangyao Hao, Mariah Weinstein, Brandon Chen, Qiuyi Liang, Qing Wang, and Dilshan Perera. Deep learning for electrocardiogram (ECG) classification. arXiv preprint arXiv:1912.03919, 2019.

[9] Anup Das, Erfan Zabeh, and Joshua Jacobs. *How to Detect and Analyze Traveling Waves in Human Intracranial EEG Oscillations?*, pages 487–505. Springer International Publishing, Cham, 2023. doi: 10.1007/978-3-031-20910-9 30. URL https://doi.org/10.1007/978-3-031-20910-9_30.

[10] Anup Das, Erfan Zabeh, Bard Ermentrout, and Joshua Jacobs. Planar, spiral, and concentric traveling waves distinguish cognitive states in human memory. bioRxiv, 2024. doi: 10.1101/2024.01.26.577456. URL https://www.biorxiv.org/content/early/2024/04/04/2024.01.26.577456.

[11] Stefano de Biase, Mariarosaria Valente, and Gian Luigi Gigli. Intractable restless legs syndrome: role of prolonged-release oxycodone–naloxone. Neuropsychiatric Disease and Treatment, 12:417, 2016.

[12] Stefano de Biase, Giulia Pellitteri, Gian Luigi Gigli, and Mariarosaria Valente. Advancing synthetic therapies for the treatment of restless legs syndrome. Expert Opinion on Pharmacotherapy, 20(16):1971–1980, 2019.

[13] Nima Ebrahimi, Hamed Moradi, Carlos Salazar Lopez, Makarand Deo, Sandeep Ramineni, and Libin Liu. A deep learning algorithm to translate and classify cardiac electrophysiology. eLife, 10:e68335, 2021. doi: 10.7554/eLife.68335.

[14] Andreas K Engel and Pascal Fries. Beta-band oscillations—signalling the status quo? Current Opinion in Neurobiology, 11(2):156–165, 2001.

[15] Andre Esteva, Alexandre Robicquet, Bharath Ramsundar, Volodymyr Kuleshov, Mark DePristo, Katherine Chou, Claire Cui, Greg Corrado, Sebastian Thrun, and Jeff Dean. A guide to deep learning in healthcare. Nature Medicine, 25(1):24–29, 2019.

[16] Zeinab Fazlali, Yadollah Ranjbar-Slamloo, Erfan Zabeh, and Ehsan Arabzadeh. Stimulation of locus coeruleus noradrenergic system modulates sensory processing and brain state in two different time scales. bioRxiv, 2023. doi: 10.1101/2020.07.09.188615. URL https://www.biorxiv.org/content/early/2023/03/07/2020.07.09.188615.

[17] Raffaele Ferri, Mauro Manconi, Debora Aricó, Chiara Sagrada, Marco Zucconi, Oliviero Bruni, Luigi Ferini-Strambi, and Stephany Fulda. Acute dopamine-agonist treatment in restless legs syndrome: effects on sleep architecture and nrem sleep instability. Sleep Medicine, 11(6):793–800, 2010.

[18] Raffaele Ferri, Filomena II Cosentino, Mauro Manconi, Francesco Rundo, Oliviero Bruni, and Marco Zucconi. Increased electroencephalographic high-frequency oscillations during the sleep onset period in patients with restless legs syndrome. Sleep, 37(8):1375–1381, 2014.

[19] Pascal Fries. Rhythms for cognition: communication through coherence. Neuron, 88(1): 220–235, 2015.

[20] Diego Garćıa-Borreguero, Richard P Allen, Ralf Kohnen, Birgit Högl, Claudia Trenkwalder, Walter Oertel, Wayne A Hening, Walter Paulus, David Rye, Arthur Walters, et al. Diagnostic standards for dopaminergic augmentation of restless legs syndrome: report from a world association of sleep medicine–international restless legs syndrome study group consensus conference at the max planck institute. Sleep Medicine, 8(5):520–530, 2007.

[21] Diego Garcia-Borreguero, James Patrick, Susan DuBrava, Philip M Becker, Andrew Lankford, Charlene Chen, John Miceli, and Lloyd Knapp. Pregabalin versus pramipexole: effects on sleep disturbance in restless legs syndrome. Sleep, 37(4):635–643, 2014.

[22] Diego Garcia-Borreguero, Michael H Silber, John W Winkelman, Birgit Högl, Judith Bainbridge, Mark Buchfuhrer, Georgios Hadjigeorgiou, Yuichi Inoue, Mauro Manconi, Wolfgang Oertel, et al. Guidelines for the first-line treatment of restless legs syndrome/willis–ekbom disease, prevention and treatment of dopaminergic augmentation: a combined task force of the irlssg, eurlssg, and the rls-foundation. Sleep Medicine, 21:1–11, 2016.

[23] Tamara Gedankien, Jennifer Kriegel, Erfan Zabeh, David McDonagh, Bradley Lega, and Joshua Jacobs. Cholinergic blockade reveals a role for human hippocampal theta in memory encoding but not retrieval. eLife, 14, 2025.

[24] Christoforos D Giannaki, Giorgos K Sakkas, Christina Karatzaferi, Georgios M Hadjigeorgiou, Efthimios Lavdas, Vassilios Liakopoulos, Yiannis Koutedakis, and Ioannis Stefanidis. Evidence of increased muscle atrophy and impaired quality of life parameters in patients with uremic restless legs syndrome. PLoS One, 6(10):e25180, 2011.

[25] Christoforos D Giannaki, Georgios M Hadjigeorgiou, Christina Karatzaferi, Maria D Maridaki, Yiannis Koutedakis, Paraskevi Founta, Nikos Tsianas, Ioannis Stefanidis, and Giorgos K Sakkas. Uremic versus idiopathic restless legs syndrome: impact on aspects related to quality of life. ASAIO Journal, 58(6):607–611, 2012.

[26] Christoforos D Giannaki, Georgios M Hadjigeorgiou, Christina Karatzaferi, Marios C Pantzaris, Ioannis Stefanidis, and Giorgos K Sakkas. Epidemiology, impact, and treatment options of restless legs syndrome in end-stage renal disease patients: an evidence-based review. Kidney International, 85(6):1275–1282, 2014.

[27] Wayne Hening, Arthur S Walters, Richard P Allen, Jacques Montplaisir, Alison Myers, and Luigi Ferini-Strambi. Impact, diagnosis and treatment of restless legs syndrome (rls) in a primary care population: the rest (rls epidemiology, symptoms, and treatment) primary care study. Sleep Medicine, 5(3):237–246, 2004.

[28] Imran H Iftikhar, Lama Alghothani, and Lynn M Trotti. Gabapentin enacarbil, pregabalin and rotigotine are equally effective in restless legs syndrome: a comparative meta-analysis. European Journal of Neurology, 24(12):1446–1456, 2017.

[29] Kasra Jalaldoust and Erfan Zabeh. A causal formulation of spike-wave duality, 2025. URL https://arxiv.org/abs/2511.06602.

[30] Stephanie R Jones. When brain rhythms aren’t ’rhythmic’: implication for their mechanisms and meaning. Current Opinion in Neurobiology, 40:72–80, 2016.

[31] Taesung Jung, Nanyu Zeng, Jason D. Fabbri, Guy Eichler, Zhe Li, Erfan Zabeh, Anup Das, Konstantin Willeke, Katie E. Wingel, Agrita Dubey, Rizwan Huq, Mohit Sharma, Yaoxing Hu, Girish Ramakrishnan, Kevin Tien, Paolo Mantovani, Abhinav Parihar, Heyu Yin, Denise Oswalt, Alexander Misdorp, Ilke Uguz, Tori Shinn, Gabrielle J. Rodriguez, Cate Nealley, Sophia Sanborn, Ian Gonzales, Michael Roukes, Jeffrey Knecht, Daniel Yoshor, Peter Canoll, Eleonora Spinazzi, Luca P. Carloni, Bijan Pesaran, Saumil Patel, Joshua Jacobs, Brett Youngerman, R. James Cotton, Andreas Tolias, and Kenneth L. Shepard. Stable, chronic in-vivo recordings from a fully wireless subdural-contained 65,536-electrode brain-computer interface device. bioRxiv, 2025. doi: 10.1101/2024.05.17.594333. URL https://www.biorxiv.org/content/early/2025/03/12/2024.05.17.594333.

[32] Konstantin Y. Kalitin, Alexey A. Nevzorov, Denis A. Babkov, Alexander A. Spasov, and Olga Y. Mukha. Deep learning analysis of intracranial eeg for recognizing drug effects and mechanisms of action, 2020. Dataset available at https://data.mendeley.com/datasets/gmkbhj28jh/1.

[33] Nancy Kopell, GB Ermentrout, Miles A Whittington, and Roger D Traub. Gamma rhythms and beta rhythms have different synchronization properties. Proceedings of the National Academy of Sciences, 97(4):1867–1872, 2000.

[34] Gaetano La Manna, Fabio Pizza, Elisa Persici, Olga Baraldi, Giorgia Comai, Maria Laura Cappuccilli, Francesca Centofanti, Elisabetta Carretta, Giuseppe Plazzi, Giuseppe Cianciolo, et al. Restless legs syndrome enhances cardiovascular risk and mortality in patients with end-stage kidney disease undergoing long-term haemodialysis treatment. Nephrology Dialysis Transplantation, 26(6):1976–1983, 2011.

[35] Anna Lindner, Krisztina Fornadi, Alpar S Lazar, Maria E Czira, Anna Dunai, Robert Zoller, Marta Visy, Andrea Kelemen, Adam Ujszaszi, Miklos Novak, et al. Periodic limb movements in sleep are associated with stroke and cardiovascular risk factors in patients with renal failure. Journal of Sleep Research, 21(3):297–307, 2012.

[36] Mikael Lundqvist, Pawel Herman, and Earl K Miller. Working memory: delay activity, yes! persistent activity? maybe not. Journal of Neuroscience, 38(32):7013–7019, 2018.

[37] Benjamin Maas, Erfan Zabeh, and Soroush Arabshahi. Quicktumornet: Fast automatic multi-class segmentation of brain tumors, 2020. URL https://arxiv.org/abs/2012.12410.

[38] Mauro Manconi, Raffaele Ferri, Marco Zucconi, Alessandro Oldani, Maria Livia Fantini, Valentina Castronovo, and Luigi Ferini-Strambi. First night efficacy of pramipexole in restless legs syndrome and periodic leg movements. Sleep Medicine, 8(5):491–497, 2007.

[39] Maya M McGregor and Alexandra B Nelson. Circuit mechanisms of parkinson’s disease. Neuron, 101(6):1042–1056, 2019.

[40] Jacques Montplaisir, Sylvie Boucher, Andre Gosselin, Guy Poirier, and Gilles Lavigne. Persistence of repetitive eeg arousals (k-alpha complexes) in rls patients treated with ldopa. Sleep, 19(3):196–199, 1996.

[41] Kavita Nagandla and Sanjay De. Restless legs syndrome: pathophysiology and modern management. Postgraduate Medical Journal, 89(1053):402–410, 2013.

[42] Maurice M Ohayon, Ruth O’Hara, and Michael V Vitiello. Epidemiology of restless legs syndrome: a synthesis of the literature. Sleep Medicine Reviews, 16(4):283–295, 2012.

[43] William G Ondo. Restless legs syndrome: pathophysiology and treatment. Current Treatment Options in Neurology, 16(11):317, 2014.

[44] Mena Raouf, Thomas J Atkinson, Michael W Crumb, and Jeffrey Fudin. Rational dosing of gabapentin and pregabalin in chronic kidney disease. Journal of Pain Research, 10:275, 2017.

[45] Robert Schmidt, Maria H Ruiz, Bjørg E Kilavik, Mikael Lundqvist, Philip A Starr, and Adam R Aron. Beta oscillations following performance feedback predict subsequent recall of task-relevant information. Scientific Reports, 10:15027, 2020. doi: 10.1038/s41598-020-72128-x.

[46] Barbara Schormair, Juliane Plag, Martina Kaffe, Nicole Groß, Darina Czamara, Walter Samtleben, Peter Lichtner, Andreas Strohle, Ioannis Stefanidis, Athanasios Vainas, et al. Meis1 and btbd9: genetic association with restless leg syndrome in end stage renal disease. Journal of Medical Genetics, 48(7):462–466, 2011.

[47] Joanna S Thurlow, Maulin Joshi, Guofen Yan, Keith C Norris, Lawrence Y Agodoa, Claudia M Yuan, and Robert Nee. Global epidemiology of end-stage kidney disease and disparities in kidney replacement therapy. American Journal of Nephrology, 52(2):98–107, 2021.

[48] Yibo Wang, Samuel H Bryant, Taosheng Cheng, Jingwei Wang, Asta Gindulyte, Benjamin A Shoemaker, Paul A Thiessen, Siqian He, and Jian Zhang. Deep learning for drug discovery and development: a review. JAMA Network Open, 3(5):e204686, 2020.

[49] Juliane Winkelmann, Alexander Stautner, Walter Samtleben, and Claudia Trenkwalder. Long-term course of restless legs syndrome in dialysis patients after kidney transplantation. Movement Disorders, 17(5):1072–1076, 2002.

[50] Erfan Zabeh, Nicholas C Foley, Joshua Jacobs, and Jacqueline P Gottlieb. Beta traveling waves in monkey frontal and parietal areas encode recent reward history. Nature Communications, 14(1):5428, 2023.

